# Multivariate analysis reveals environmental and genetic determinants of element covariation in the maize grain ionome

**DOI:** 10.1101/241380

**Authors:** Alexandra Asaro, Brian P. Dilkes, Ivan Baxter

## Abstract

Plants obtain elements from the soil through genetic and biochemical pathways responsive to physiological state and environment. Most perturbations affect multiple elements which leads the *ionome*, the full complement of mineral nutrients in an organism, to vary as an integrated network rather than a set of distinct single elements. To examine the genetic basis of covariation in the accumulation of multiple elements, we analyzed maize kernel ionomes from Intermated B73 × Mo17 (IBM) recombinant inbred populations grown in 10 environments. We compared quantitative trait loci (QTL) determining single-element variation to QTL that predict variation in principal components (PCs) of multiple-element covariance. Single-element and multivariate approaches detected partially overlapping sets of loci. In addition to loci co-localizing with single-element QTL, multivariate traits within environments were controlled by loci with significant multi-element effects not detectable using single-element traits. Gene-by-environment interactions underlying multiple-element covariance were identified through QTL analyses of principal component models of ionome variation. In addition to interactive effects, growth environment had a profound effect on the elemental profiles and multi-element phenotypes were significantly correlated with specific environmental variables.

**Author Summary:** A multivariate approach to the analysis of element accumulation in the maize kernel shows that elements are not regulated independently. By describing relationships between element accumulation we identified new genetic loci invisible to single-element approaches. The mathematical combinations of elements distinguish groups of plants based on environment, demonstrating that observed variation derives from interactions between genetically controlled factors and environmental variables. These results suggest that successful application of ionomics to improve human nutrition and plant productivity requires simultaneous consideration of multiple-element effects and variation of such effects in response to environment.

## Introduction

Elements are distinct chemical species, and studies of element accumulation frequently investigate each element separately. There is overwhelming evidence, however, that element accumulations covary due to physical, physiological, genetic, and environmental factors. In a dramatic example in *Arabidopsis thaliana*, a suite of elements responds to Fe deficiency in such a concerted manner that they can be used to predict the deficiency or sufficiency of Fe for the plant more accurately than the measured level of Fe in plant tissues [1]. The basis of this covariation can be as simple as co-transport of multiple elements. IRT1 is a metal transporter capable of transporting Fe, Zn, and Mn. IRT1 is upregulated in low Fe conditions resulting in an environmentally-dependent link between Fe and other ions [2]. Other pairs of co-regulated elements, such as Ca and Mg, which have been shown to exhibit shared genetic regulatory networks in *Brassica oleracea* [3], should be affected identically, or predictably, by genetic variation. When *A. thaliana* recombinant inbred line populations were grown in multiple environments, genetic correlations among Li-Na, Mg-Ca, and Cu-Zn were observed across all environments while Ca-Fe and Mg-Fe were only correlated in a subset of environments [4]. Shared genetic control of ion transport without substantial environmental responsiveness should result in the former pattern, along with significantly less capacity for homeostasis across environmental concentrations and availabilities of elements. Environmentally-responsive molecular mechanisms, reminiscent of *IRT1* upregulation, could result in environmentally-variable patterns of correlations. Baxter et al. previously tested element seed concentrations for correlations in the maize Intermated B73 × Mo17 (IBM) recombinant inbred population, finding several correlated element pairs, the strongest of which was between Fe and Zn [5]. Yet, few QTL impacting more than one element were found, likely due to effects on multiple elements being below the threshold of observation when mapping on single element traits with limited numbers of lines. These observations indicate that, while understanding the factors driving individual element accumulation is important, we must consider the ionome as a network of co-regulated and interacting traits [6]. We propose that formally considering this coordination between elements can provide deeper insight than focusing on each element in isolation.

Multivariate analysis techniques, such as principal components analysis (PCA), can reduce data dimension and summarize covariance of multiple traits as vectors of values by minimizing the variances of input factors to new components. When multiple phenotypes covary, as occurs for the elements in the ionome, this approach may complement single element approaches by describing trait relationships. In studies on crops such as maize, PCA has been used as a strategy to consolidate variables that may be redundant or reflective of a common state [7–9]. PCA has proved useful in previous QTL mapping efforts, facilitating detection of new PC QTL not found using univariate traits in analyses of root system architecture in rice [10] and kernel attributes, ear architecture, and enzyme activities in maize [11–13]. In the current study, we expect that elemental variables are functionally related and therefore need new traits to describe elemental covariation. Since we do not know the exact nature of these relationships, and the ionome varies depending on environment, PCA is useful in that it does not require *a priori* definition of relationships between variables. If the PCA approach leads to novel loci and insights into how the ionome is functioning, it will be a valuable addition to the study of mineral nutrient regulation.

Here we show that developing multivariate traits reveals environmental and genetic effects that are not detected using single elements as traits. We performed PCA on element profiles from the maize IBM population [14] grown in 10 different environments. Different relationships between elements were identified that depended on environment. QTL mapping using multi-element PCs as traits was carried out within each environment separately. Comparing these multivariate QTL mapping results to previous QTL analyses of the same data using each single element as traits for QTL analysis [15] demonstrates that a multivariate approach uncovers unique loci affecting multi-element covariance. Additionally, an experiment-wide PCA performed on combined data from all environments produced components capable of separating lines by environment based on their whole-ionome profile. These experiment-wide factors, while representative of environmental variation, also exhibited a genetic component, as loci affecting these traits were detected through QTL mapping.

## Results

### Summary of Data Collection and Previous Analysis of Single Element Traits

We previously acquired data on 20 elements measured in the seeds from *Zea mays* L. Intermated B73 × Mo17 recombinant inbred line (IBM) populations [14] grown in 10 different location/year settings [15]. This work is briefly summarized here as it serves as the basis of our comparison. The kernels came from RILs of the IBM population cultivated across six locations and five years. Quantification of the accumulation of 20 elements in kernels was done using inductively coupled plasma mass spectrometry (ICP-MS). Weight-adjusted element measurements were used for a QTL analysis to detect loci contributing to variation in seed element contents [15]. The current study is motivated by previous demonstrations of elemental correlations and mutant phenotype analyses which indicate extensive relationships between elements [1, 4]. To explore this formally, we further analyzed these data from a multiple-element perspective.

### Element to Element Correlations

Several elements were highly correlated across the dataset, exhibiting pairwise relationships among lines in a given environment that passed a conservative Bonferroni correction for multiple tests. We detected 209 pairs of elements that were genetically correlated out of 1,900 possible correlations across environments (190 pairs per environment). We expect that evidence of robust genetic control would be provided by repeated observation of trait correlations in multiple environments. Of the six locations included in this experiment, we obtained data from three locations (FL, IN, and NY) from plant material grown in two different years. Seven element-pairs were correlated in five or more of these six environments: Mn and Mg, Ca and Sr, S and P, K and P, P and Mg, S and Mg, and Fe and Zn (Fig 1). Other element-pair correlations were driven by the genetic variation of the IBM in fewer environments. For example, Mn and P were correlated in FL05, NY05, and NY12 (r_p_ = 0.50, 0.48, 0.51) but were not significantly correlated in FL06, IN09, or IN10 (r_p_ = 0.31, 0.20, 0.18). Thus, while some correlations exist in multiple years and multiple locations, element correlations were affected by both location and year.

**Fig 1.**
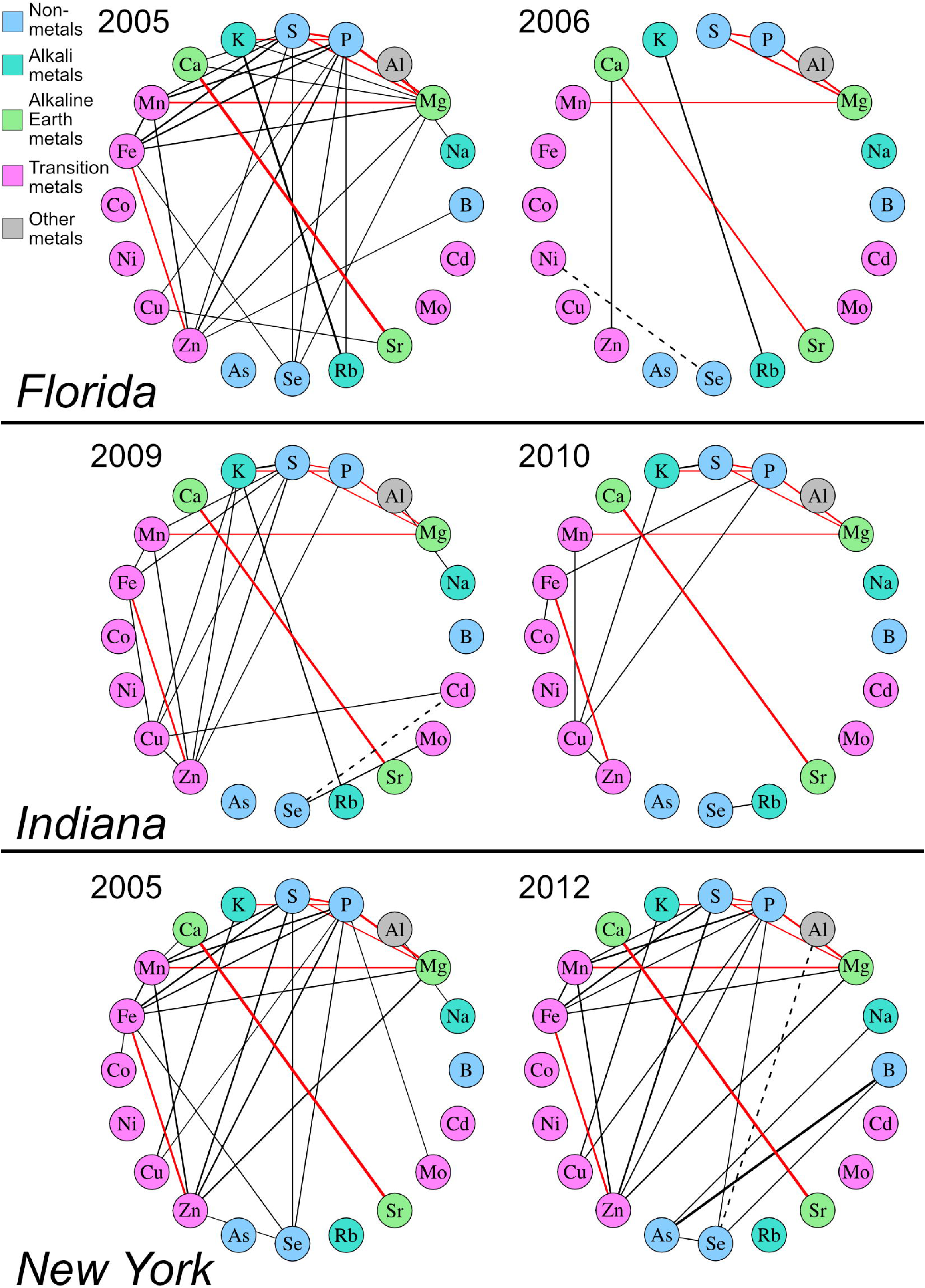
Element Correlations Diagrams for Locations with Repeated Measurements. Pairwise correlations of 20 kernel elements in varying environments, shown for the experiments within locations having data from multiple years (FL, IN, and NY). Correlations were calculated as the Pearson correlation coefficient (r_p_) between concentration values for each element pair. Significance was evaluated using a Bonferroni correction for multiple tests within each environment and set at a corrected p value of 0.05. Lines between elements represent significant pairwise correlations, weighted by strength of correlation. Positive and negative correlations are represented as solid and dashed lines, respectively. Red lines indicate correlations present in at least 5 of the 6 environments shown.

In our previous single-element QTL analysis of these data, loci comprising QTL for two or more different elements were detected (Table 1). This shared genetic control of multiple elements was readily apparent in the trait correlations calculated within environments, as five of the nine shared-element QTL exhibited corresponding element pair correlations within the given environment. For example, phosphorous, which was in three of the seven most reproducible element-pair correlations, exhibited the highest incidence of shared QTL with other elements. These included shared QTL between P accumulation and all three of the reproducibly P-correlated elements: S and the cations K and Mg. In addition, P was affected by the only QTL shared between more than two elements, which affected P, S, Fe, Mn, and Zn accumulation in NY05 (Fig 2). Consistent with the possibility of variation in transport processes affecting element accumulation correlations, shared QTL were frequently found between elements with similar structure, charge, and/or type, such as Ca and Sr or Fe and Zn. These element correlations and post-hoc comparisons of shared QTL localizations suggest a genetic basis for covariance of the ionome in the RIL population.

**Fig 2.**
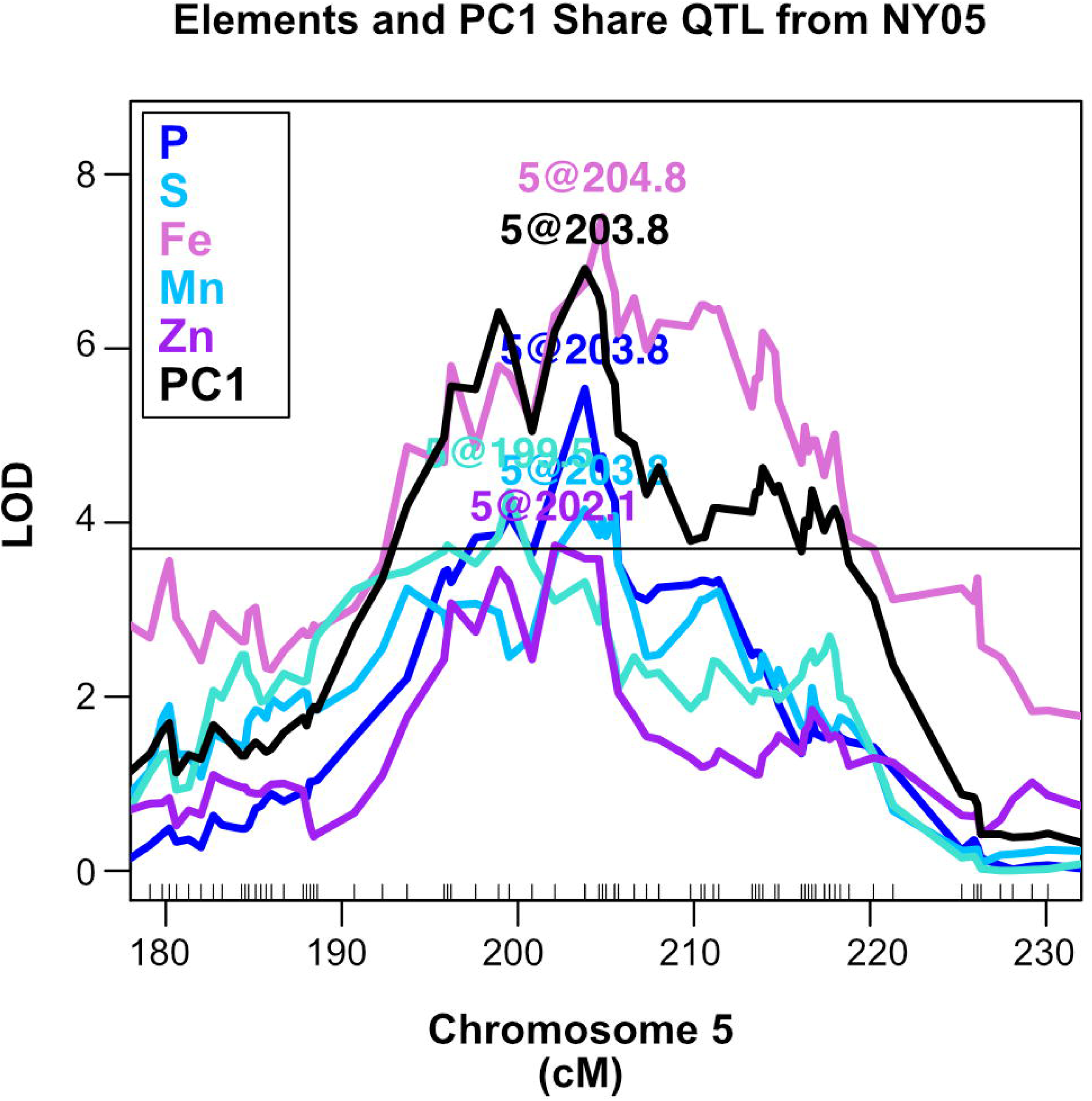
Multiple Element QTL. Stepwise QTL mapping output from the NY05 population for P, S, Fe, Mn, Zn, and PC1. Position in cM on chromosome 5 is plotted on the x-axis and LOD score is shown on the y-axis. 95^th^ percentile of highest LOD score from 1000 random permutations is indicated as horizontal line.

**Table 1.** QTL Affecting Variation for Multiple Elements in the Same Environment.

| Environment | Chr | Pos (cM)<br>† | El 1 | El 2 | El 3 | El 4 | El 5 |
| --- | --- | --- | --- | --- | --- | --- | --- |
| NY05 | 1 | 400 | Mn | Ni | --- | --- | --- |
| NY05 | 3 | 323 | Sr | Ca | --- | --- | --- |
| NY05 | 5 | 201 | Mn | Zn | P | S | Fe |
| NY06 | 1 | 532 | Mn | Mg | --- | --- | --- |
| IN09 | 4 | 306 | Fe | K | --- | --- | --- |
| IN10 | 2 | 213 | Mo | Cd | --- | --- | --- |
| NY12 | 5 | 203 | Zn | Fe | --- | --- | --- |
| FL05 | 1 | 230 | B | Mn | --- | --- | --- |
| FL05 | 4 | 159 | Fe | Zn | --- | --- | --- |

†Average position

### Principle Components Analysis of Covariance for Elements in the Ionome

To better describe multi-element correlations and thereby detect loci controlling accumulation of two or more elements, we derived summary values representing the covariation of several elements. We implemented an undirected multivariate technique, principal components analysis (PCA), for this purpose. PCA reduced correlated elements into principal components (PCs), orthogonal variables that account for variation in the original dataset, each having an associated set of rotations (also known as loadings) from the input variables. After removing elements prone to analytical artifacts, PCA was conducted using the remaining 16 elements from each of the 10 environments separately. This produced 16 principal components in each environment (S1 Fig) of which we retained for further analysis only PCs representing more than 2% of the total variation. This resulted in as few as 11 and as many as 15 PCs depending on environment.

Remarkably, there is substantial overlap in the loadings of many elements in the first and second PCs across some environments, suggesting a reproducible effect of genetic variation on the ionome in these environments (Fig 3). Additionally, the loadings of elements are consistent with the pair-wise relationships observed in the element-by-element correlations. For example, Ca and Sr frequently load PCs in a similar direction. The PC loadings derive from inputs of several elements to a single PC variable. All retained PCs in all 10 environments have a loading contribution of at least .25 in magnitude from two or more elements. While some patterns existed across environments, many PC loadings differed in both magnitude and direction according to environment, suggesting instability of element-pair correlations across the environments. As these PCs were separately calculated in each environment, we compared PC traits from different environments. We used correlation tests of element loadings in PCs to identify PCs from different environments that were constructed from similar relationships. Because loading direction is arbitrary, both strong positive and strong negative correlations were examined. 52 pairs of PCs exhibited loadings correlations with a Pearson correlation coefficient greater than 0.75 or less than −0.75 (S2 Fig). Thus, the PC analyses produced pairs of correlated PCs in different locations that, while not necessarily recovered in the same order, derived from similar patterns of elemental variation.

**Fig 3.**
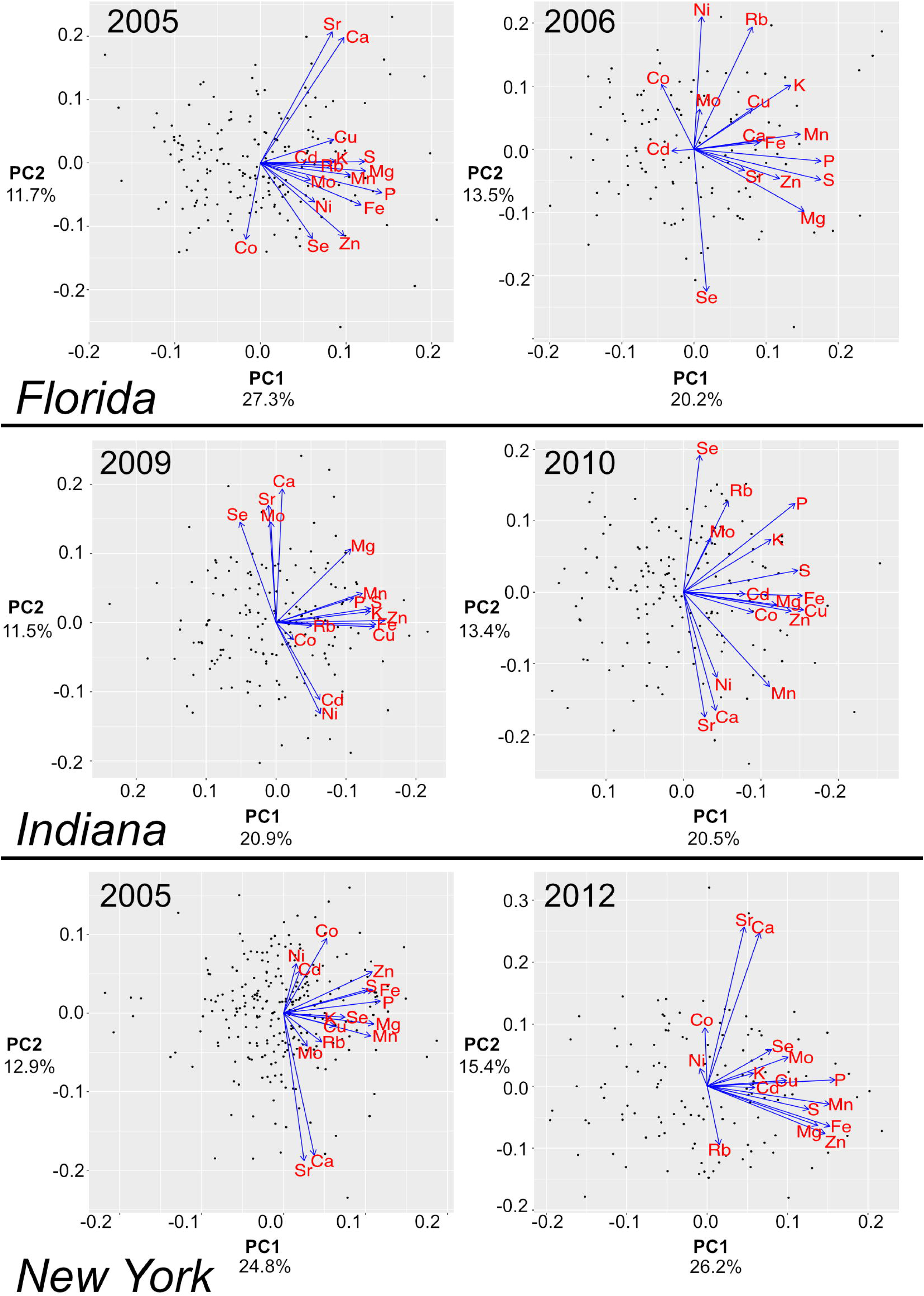
PCA Plots in Multiple Environments. PCA plots showing PC1 and PC2 loadings in different years in three locations (FL, IN, and NY). PC1 and PC2 values for each line are plotted as points and PC1 and PC2 loadings of each element are indicated by blue arrows. The data for different years for each of three locations, FL, IN, and NY are plotted. The percent of total variation explained by each PC is labeled on the axes.

### QTL Mapping of Ionomic Covariance Components

The PCs from each environment were used as traits for QTL detection. Stepwise QTL mapping using these derived traits yielded 93 QTL that exceeded an estimated statistical threshold of α=0.05 from within-environment permutations (Fig 4C). 56 of these QTL affecting multiple-element covariance components overlapped with previously detected single-element QTL in the same environment [15] (Fig 4A). In some cases, two or more PC traits within an environment resolved to one single-element QTL. This was observed particularly for elements with strong effect QTL, such as Mo, Cd, and Ni. For example, in IN10, PC2 and PC10 both have QTL that co-localize with the Cd QTL on chromosome 2. Likewise, in NY05, PC3, PC5, PC6, and PC9 all detect a QTL coinciding with the large-effect Ni QTL on chromosome 9. These PCs within a single environment all have varying levels of Ni contribution, as well as varying levels of contribution from other elements. Although the relationship among elements described by each PC is distinct, the same single-element locus can be detected due to that locus affecting an element that is present within each set of relationships. This repeated detection of the same locations contributes to the higher number and proportion of detected PC QTL that were shared with element QTL (56/93) than element QTL that were shared with PC QTL (18/79), although the same genomic locations underlie this overlap. 37 PC QTL were detected at loci not seen using single element traits, demonstrating that PC traits can outperform single element data for the detection of shared genetic control of correlated characters. For instance, two PC5 QTL from the NY06 growout were located on chromosome 1 at positions distinct from any elemental QTL (Fig 4B). QTL mapping on single elements may not have the power to detect loci with small coordinate effects on several elements. So as to not inflate PC-specific QTL, they are defined here as QTL greater than 25 cM away from any elemental QTL in the same environment.

**Fig 4.**
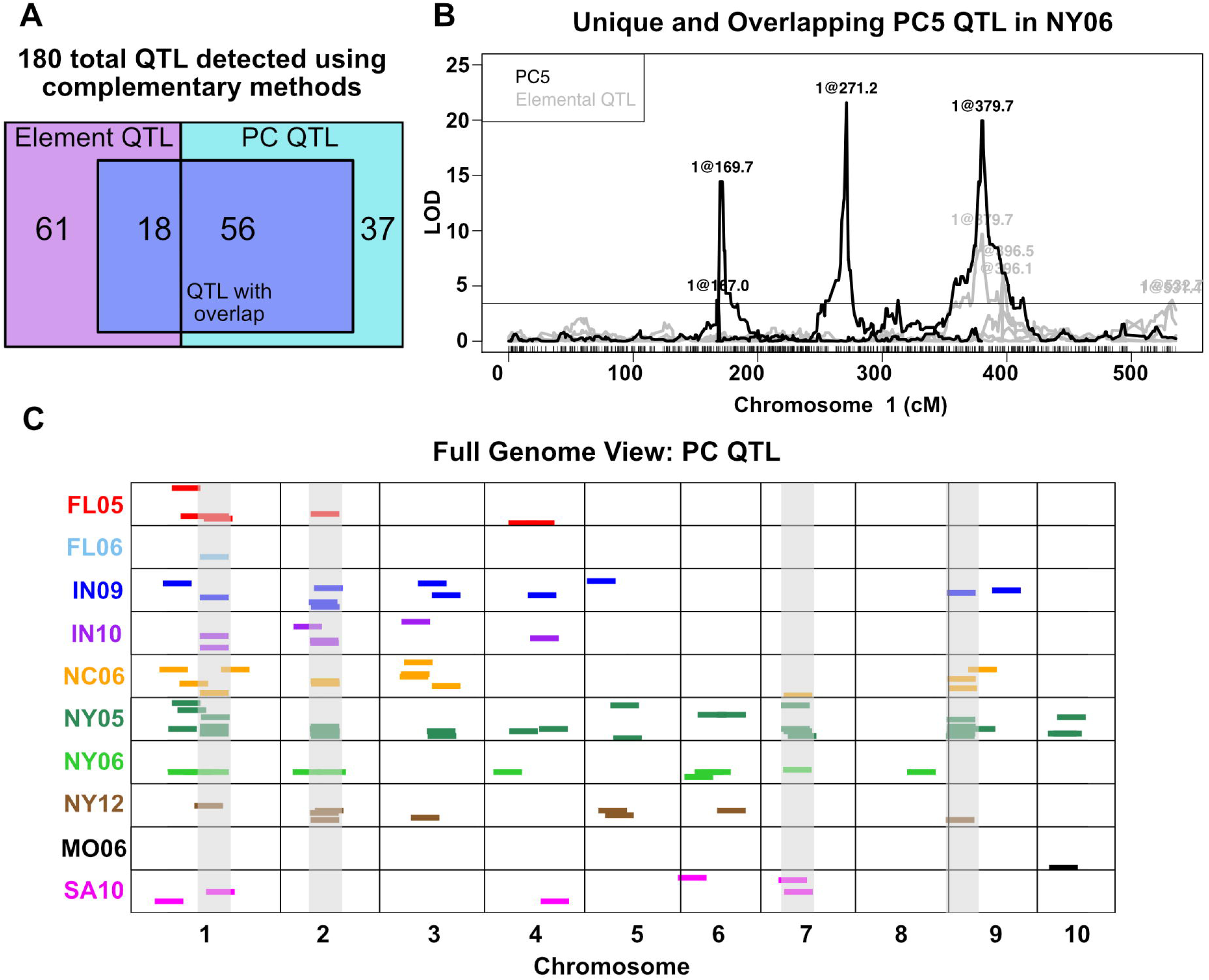
Principal Component QTL from 10 environments. PCs were derived from elemental data separately in each of 10 environments and used as traits for QTL mapping. (A) 172 total element and PC QTL were mapped. The two boxes represent the 79 and 93 elemental and PC QTL, respectively. 18 element QTL overlap with PC QTL from the same environment. S6 PC QTL overlap with element QTL from the same environment. Sets of non-unique QTL are shown in the center box. QTL unique to elements, 61, and to PCs, 37, are shown outside of the shared box. (B) QTL mapping output for PCS from the NY06 population. Position on chromosome 1 is shown on the x-axis, LOD score is on the y-axis. All significant NY06 element QTL on chromosome 1 are shown in grey (α = 0.05). Two PCS QTL, at 169.7 and 271.2 cM, are unique to PCS and do not overlap with any elemental QTL. A PCS QTL at 379.7 cM is shared with a molybdenum QTL. (C) Significant PC QTL (α = 0.05) for PCs in 10 environments. QTL location is shown across the 10 chromosomes on the x-axis. Environment in which QTL was found is designated by color. QTL are represented as dashes of uniform size for visibility. Four regions highlighted in grey represent the four loci found for multiple PC traits in multiple environments (> 2).

PC QTL analysis captured previously observed single-element QTL shared between elements within a particular environment. Of the nine loci affecting variation for multiple elements in the same environment (Table 1), four loci also impact variation for a PC trait in that environment (Table 2). For example, in NY05, a QTL for PC1 overlaps the QTL that was detected in the single element analyses of P, S, Fe, Mn, and Zn on chromosome 5 (Fig 2). The PC QTL in this case was as strong as the association between the locus and Fe accumulation and more significant than the P, S, Mn, and Zn elemental QTL. Thus, QTL mapping a multi-element PC was as strong as the best single-element approach for previously detected QTL. For traits that cause variation in multiple elements, such as root structure, the PC approach may be preferable to single elements, particularly in cases where single element changes are of small effect or below detection limits while concerted changes to multiple elements display a larger effect.

**Table 2.**
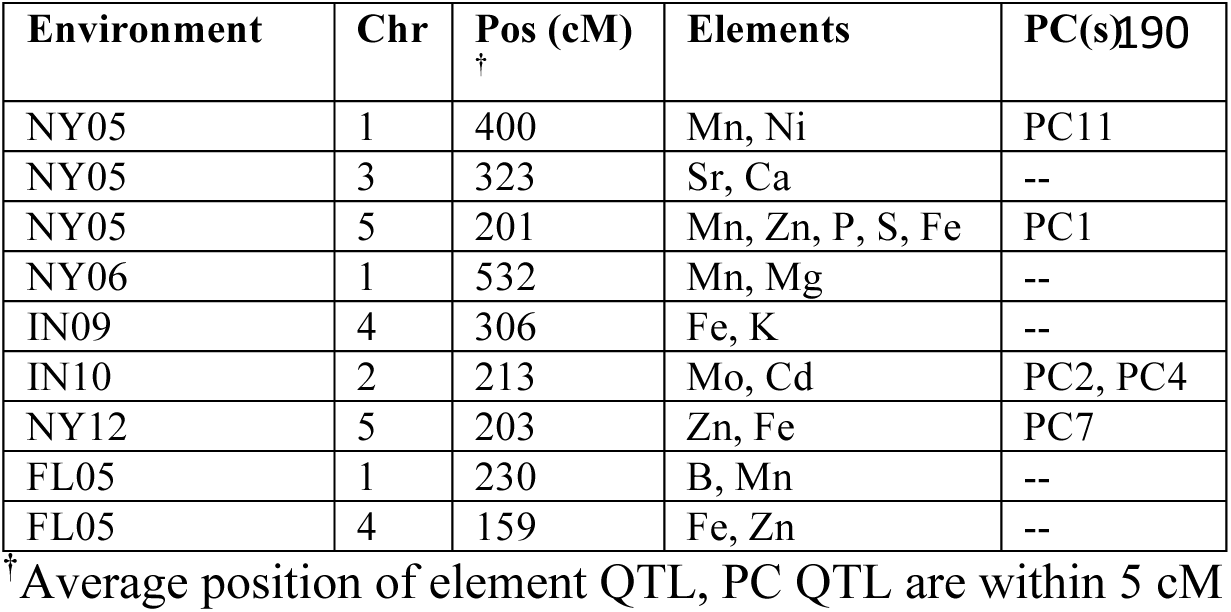
QTL for Multiple Elements and PC(s) in the Same Environment.

| Environment | Chr | Pos (cM)<br>† | Elements | PC(s) |
| --- | --- | --- | --- | --- |
| NY05 | 1 | 400 | Mn, Ni | PC11 |
| NY05 | 3 | 323 | Sr, Ca | -- |
| NY05 | 5 | 201 | Mn, Zn, P, S, Fe | PC1 |
| NY06 | 1 | 532 | Mn, Mg | -- |
| IN09 | 4 | 306 | Fe, K | -- |
| IN10 | 2 | 213 | Mo, Cd | PC2, PC4 |
| NY12 | 5 | 203 | Zn, Fe | PC7 |
| FL05 | 1 | 230 | B, Mn | -- |
| FL05 | 4 | 159 | Fe, Zn | -- |

†Average position of element QTL, PC QTL are within 5 cM

We compared PCs from different environments and looked for overlapping QTL among PCs in different environments with correlated loadings. Of the 52 PC pairs with correlated loadings, 37 had no QTL for one or both of the PCs, consistent with a shared environmental factor variable in those fields as the basis of that variation. Of the remaining 15 pairs with at least one QTL detected for each member of the pair, PCs in five pairs had shared QTL. In all five cases, the QTL shared between these pairs of PCs correspond to a large-effect single-element QTL. Six PC traits belonging to three correlated pairs, PC4 in NY05 and PC6 in IN09 (r_p_ = 0.81), PC4 in FL05 and PC3 in NY05 (rp = −0.84), and PC3 in IN10 and PC2 in NC06 (rp = 0.89), detected a QTL coinciding with a Mo QTL, a locus on chromosome 1 encoding the ortholog of the *A. thaliana* MOT1 molybdenum transporter. The same scenario exists for PC2 in IN09 and PC2 in NY05 (r_p_ = −0.78), both affected by the QTL on chromosome 2 that had a strong effect on Cd in our single-element QTL mapping experiments. Finally, PC8 in NC06 and PC5 in NY05 (r_p_ = 0.76) both map to a large-effect Ni QTL. Despite the resolution to QTL detected in a single-element analysis, in all of these cases correlations between loadings were not driven by a single element, but rather by similar loadings for most elements (S2 Fig). In addition to overlaps at these strong-effect single element QTL, 6 other pairs of correlated PCs have QTL that do not overlap. Correlated PCs with QTL at different chromosomal positions in different environments could be due to states, such as increased root system volume or iron deficiency, that may arise from distinct processes in each environment yet can generate a consistent physiological response. In these cases, the ionome displays similar trait covariance but different genetic architecture consistent with genotype by environment interactions.

The PC approach also detected a QTL that was found for different single elements depending on environment. The same locus on chromosome 7 encoded QTL for three different elements, Cu, K, and Rb, each in a different environment. K and Rb are chemical analogs. Failure to detect this QTL as affecting both elements in the same environment may simply indicate the poor power to detect all QTL, resulting in false negative results. It is less likely, but possible, to result from incorrect assessment of a shared genetic basis due to fortuitous linkage of multiple loci. Using the PC traits, we detected QTL at this position in these same three environments and a fourth environment. Thus, PCs can provide an improved estimate for the genetic effect on phenotypic variance for multi-element traits. In SA10, no QTL were mapped for Cu, Rb, or K alone. Yet, this locus was detected as significantly affecting variation in PC9 calculated from SA10, the loadings of which show a strong contribution from Cu and Rb.

The identification of both unique and previously observed QTL through this multivariate approach demonstrates the complementary nature of working with trait covariance as well as the component traits and supports previous work showing that elemental traits are mechanistically interrelated. The repeated finding of results consistent with GxE led us to investigate this formally.

### QTL by Environment Interactions

Our prior analyses found QTL by environment interactions contributing to accumulation of single elements [15]. Given element correlations and partially overlapping sets of element and PC QTL, we expect to detect QTL by environment interactions that impact multi-element traits. To look at the effects of environment on genetic regulation of multi-element phenotypes, we conducted another PCA, this time on element concentrations of lines from all environments combined. If the genetic and environmental variances do not interact, we expect some PCs will reflect environmental variance and others will reflect genetic variance. However, if the ionome is reporting on a summation of physiological status that results from genetic and environmental influences, some PCs calculated from ionomic traits should be both correlated with environmental factors and result in detectable QTL.

#### PCA across environments

The covariance between element accumulation data across all environments was summarized using principal components analysis. Elements prone to analytical artifacts (B, Na, Al, As) were removed prior to analysis. 16 across-environment PCs (aPCs) describing the covariation of the ionome were calculated for every RIL in every environment.

Out of a concern that the different lines present in each growout unduly influenced the construction of PCs specific to each environment, we performed the following tests. First, we looked at only those locations where two or more growouts were performed, so that location replication might be considered. Second, to identify a balanced sample set present in all environments, we identified the lines that were grown in all of these six growouts. PCA of the 16 element measurements was conducted across environments (S3 Fig) and the loadings of each element into each PC were recorded. Thus, the loadings of the 16 elements in the PCA were calculated from a set of common genotypic checks distributed within each environment. We used these loadings to calculate PCA projections (PJs) from all lines in all environments. In this way we made comparisons of the same calculated values in each environment. We found that the PJs and aPCs were strongly correlated; PJ1 and aPC1 were nearly identical (r_p_ = .998) and PJs 2–5 correlated with at least one of aPCs 2–5 at r_p_ > .66. The correlations between the loadings from PJs and aPCs reflected these same patterns. To reduce the incidence of artifacts or overfitting, aPCs accounting for less than 2% of the total variation were eliminated for further analyses, leaving seven aPCs.

Growth environment had a significant effect on all aPCs (p < 0.001). The first two aPCs were highly responsive to the environment (Fig 5). The lines from each environment cluster together when plotting aPC1 vs aPC2 values, with distinct separation between environments and years. In order to identify environmental factors responsible for ionome covariance, weather station and soil data from all environments except SA06 were recovered from databases (see methods). Correlations were calculated between season-long or quarter-length summaries of temperature and the aPC values for the nine environments. The weather variables, all temperature-based, were not correlated with aPCs in many cases, although correlations exceeding r_p_ = 0.50 were observed for aPCs 2,4, and 5 (Fig 6A). The strongest correlation observed for aPC1 was with average maximum temperature in the fourth quarter of the growing season (r_p_ = 0.35) (Fig 6B) while the highest observed for aPC2 was for average maximum temperature during the third quarter (r_p_ = 0.58) (Fig 6C). The relatively small number of environments, substantial non-independence of the weather variables, and likely contribution of factors other than temperature limit the descriptive power of these correlations.

**Fig 5.**
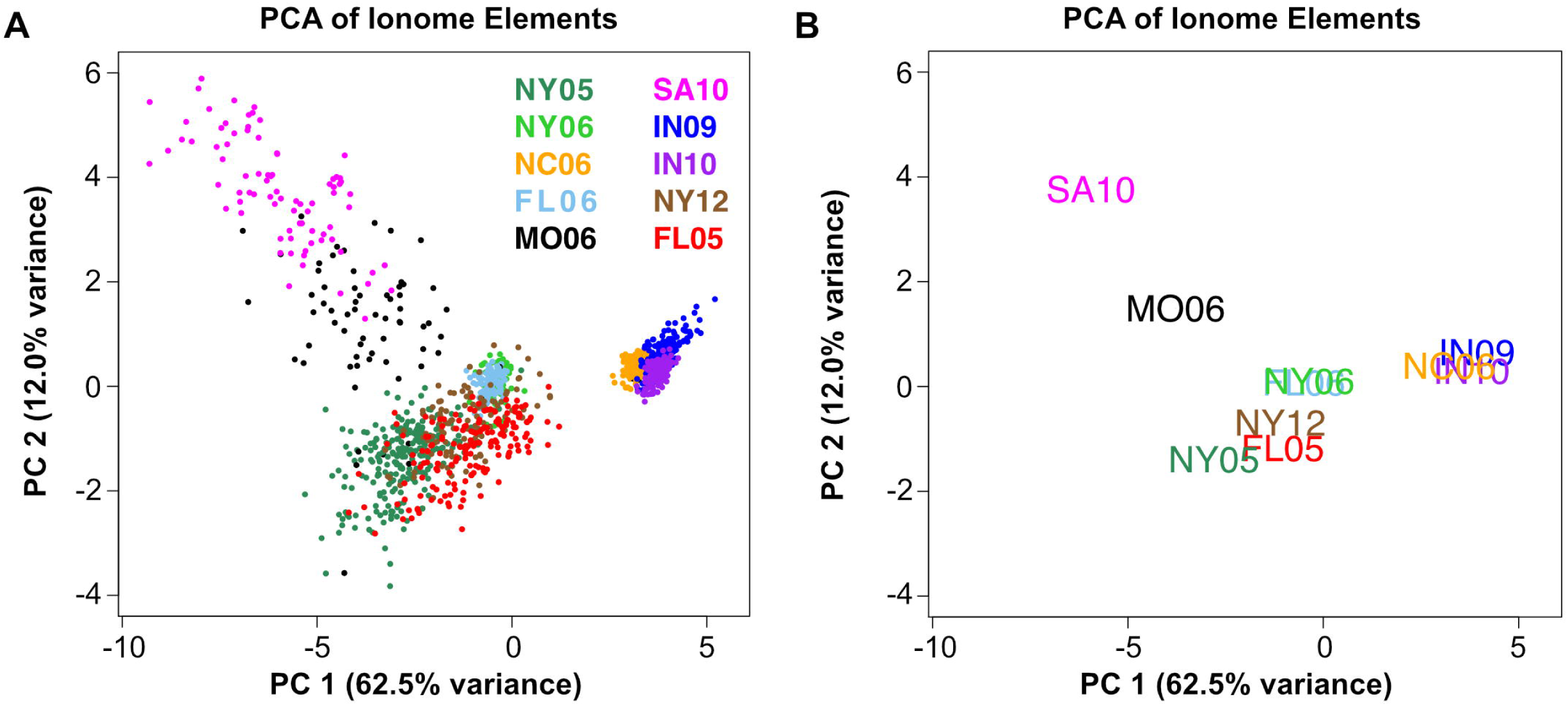
PCA Separates Lines by Environment. PC1 and PC2 separate lines by environment. Points correspond to lines, colored by their environment. (A) Across-environment PC1 vs PC2 values for each line, colored by environment. Percentage of total variance accounted for by each PC indicated on the axes. (B) Average across-environment PC1 vs PC2 values for all lines in each environment.

The lack of particularly strong correlations between the first two aPCs and temperature variables suggests that other variables, possibly field to field variation in soil composition, fertilizer application, humidity, or abiotic factors, are likely to have an influence. Correlations were also calculated between environment averages of the PCs and soil variables (Fig 6D). While the majority of these features were not found to be highly correlated with aPCs, we did observe a strong negative correlation between aPC2 and soil pH (r_p_ = −.78) (Fig 6E).

**Fig 6.**
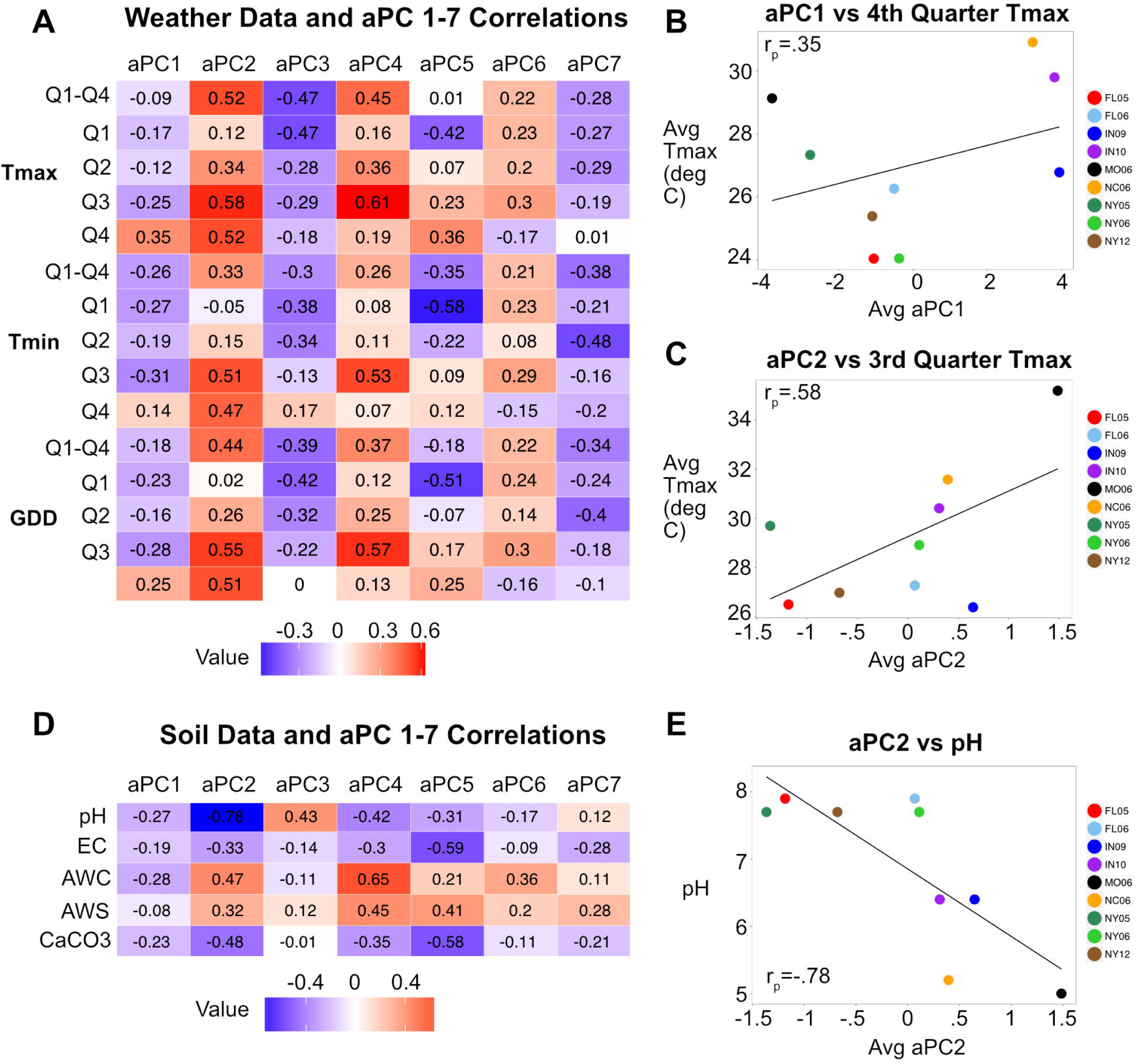
aPC and Weather Variable Correlations. (A) Heatmap showing Pearson correlation coefficients (r_p_) between averaged aPC 1–7 values across environments and averages for maximum temperature, minimum temperature, and GDD across the growth season and for each quarter of the season. Red and blue intensities indicate strength of positive and negative correlations, respectively. (B) Average aPC1 values for 9 environments vs. average maximum temperature for each environment over the fourth quarter of the growing season. Points colored by environment. Pearson correlation coefficient is shown within the graph. (C) Average aPC2 values for nine environments vs. average maximum temperature for each environment over the 3rd quarter of the growing season. (D) Heatmap showing correlations between aPCs 1–7 and soil attributes: pH, electrical conductivity (EC), available water capacity (AWC), available water storage (AWS), and calcium carbonate (CaCO3). (E) Average aPC2 values vs. pH.

In order to determine genetic effects on these components, the calculated values for aPC1 through aPC7 were used as traits for QTL analysis in each of the 10 environments. Unlike the earlier described PCAs done in environments separately, these aPCs are calculated across all environments and are therefore comparable between environments. QTL mapping detected at least four loci controlling each aPC and a total of 38 QTL. Nine of these QTL were found in common across multiple environments and 29 were only detected in a single environment (Fig 7). Of the aPC QTL, the highest LOD score QTL were present in multiple environments and corresponded to the locations of the two strongest single element QTL previously detected from the same data (Mo on chromosome 1 and Cd on chromosome 2). The detection of QTL, together with the strong environmental determination of aPCs 1–7, demonstrates that ionomic covariation results from coordinate environmental and genetic variation.

**Fig 7.**
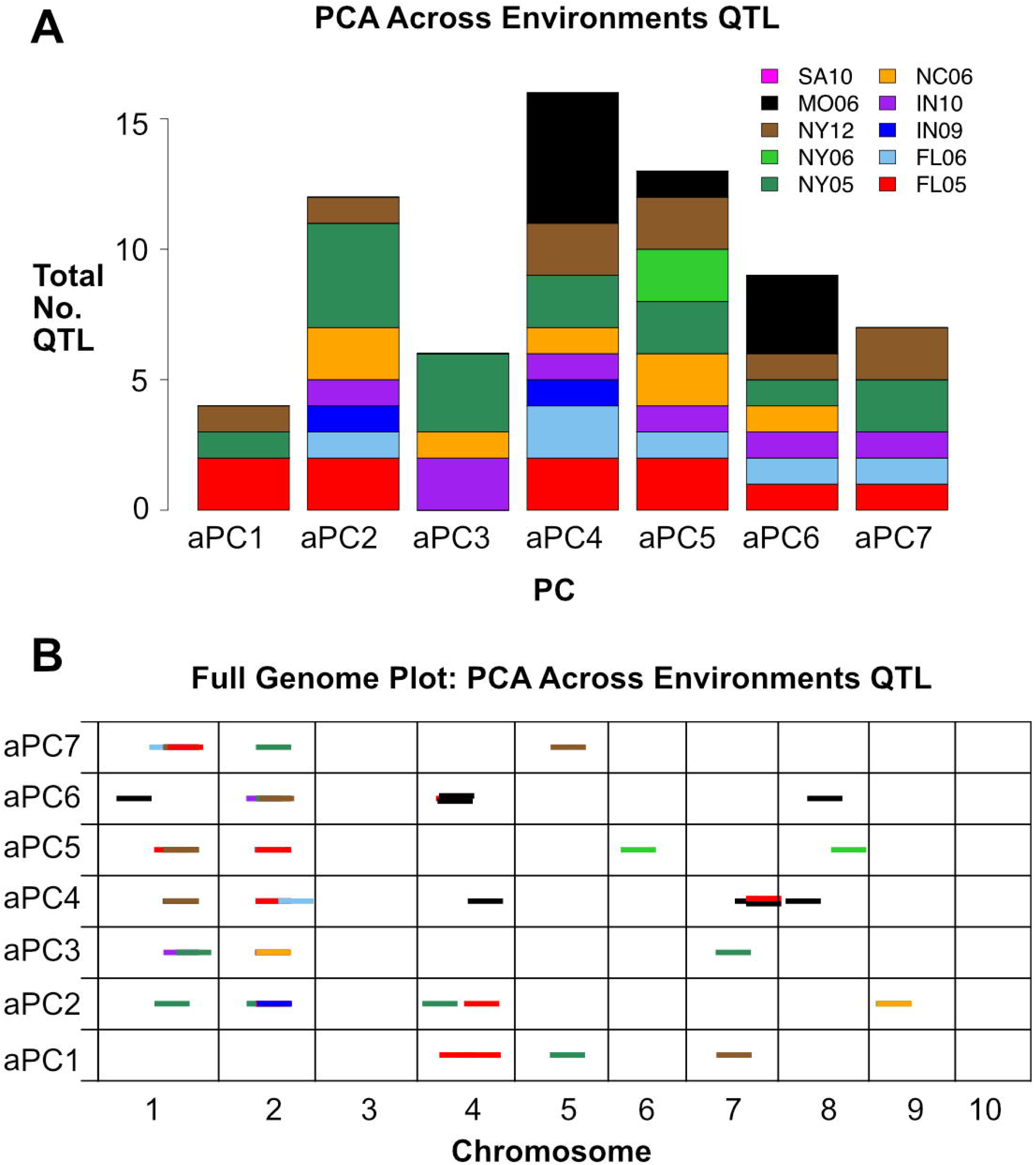
Across-Environment PCA QTL in 10 Environments. QTL identified for across environment PCA traits (aPCs 1–7). (A) Total number of QTL detected for each aPC, colored by environment. (B) Significant QTL (α = 0.0S) for aPCs 1–7. QTL location is shown across 10 chromosomes (in cM) on the x-axis. Dashes indicate QTL, with environment in which QTL was found designated by color. All dashes are the same length for visibility.

Based on the stochastic detection of QTL in only a subset of growth environments, substantial interaction between the environment aPC QTL is expected. A QTL of particular interest is the aPC2 QTL detected for Mo at the ortholog of the *MOT1* locus. Previous studies have demonstrated a connection between pH and molybdenum, with Mo availability in soil being increased by high pH. It was found that the *MOT1* locus in *A. thaliana* determines response to pH changes and resultant changes in Mo availability in an allele-specific manner, suggesting an adaptive role for variation in *MOT1* with respect to soil pH [16]. The correlation between aPC2 and pH was significant and aPC2 identified a QTL coinciding with a Mo QTL suggesting genetic variation in pH-dependent changes to Mo availability across environments. The loading magnitude for Mo into aPC2 is 0.21 but Co, Ni, Rb, and Cd contribute even more, with loading magnitudes of 0.24, 0.46, 0.55, and 0.41, respectively. QTL for aPC2 also overlap with QTL for Cd and Ni. With aPC2 representing several elements, the correlation with soil pH and overlap with single element QTL may reflect a multi-element phenotype responding to changes in pH. Further investigation is needed to molecularly identify the genes underlying aPC QTL, their biological roles, and their interaction with specific environmental variables.

## Discussion

In this study, we demonstrate that multi-trait analysis is a valuable approach for understanding the ionome. The ionome is a homeostatic system, and effects on one element can affect other elements [1]. Many biological processes in maize have the potential to impact several elements. Indirect effects on a suite of elements have been demonstrated for numerous physiological states. Radial transport of nutrients is controlled in part by endodermal suberin, the structure and deposition of which can adapt in a highly plastic manner in response to deficiencies in K, S, Na, Fe, Zn, and Mn, potentially modifying transport of additional elements [17]. Other examples of indirect effects can be found in Arabidopsis *TSC10A* mutants with reduced 3-ketodihydrosphinganine (3-KDS) reductase activity. Because 3-KDS reductase is needed for synthesis of the sphingolipids that regulate ion transport through root membranes, these mutants exhibit a completely root-dependent leaf ionome phenotype of increased Na, K, and Rb, and decreased Mg, Ca, Fe, and Mo [18].

In line with the abundance of concerted element changes seen in ionome mutants, we detected elemental correlations and QTL that were present for more than one element. Phosphorous exhibited the greatest number of QTL overlap with other elements, including the cations K and Mg. Phosphorous is a central nutrient in plant development and regulates other elements, complexing with cations in the form of phytic acid in maize seeds [19]. Additional shared QTL included those between Ca and Sr, Mo and Mn, and Zn and Fe. Ca and Sr are chemical analogs while Zn and Fe regulation have been linked at the physiological and molecular level [6, 20]. Mo and Mn have roles in protein assimilation and nitrate regulation [21, 22] and exhibit a regulatory relationship [23]. Thus, these shared QTL likely reflect genetic polymorphisms affecting the activity of multi-element regulatory genes or genetic changes targeted to a single element with pleiotropic effects on other elements via homeostatic mechanisms.

The 37 PC-specific loci identify novel loci in maize with the potential to expand our understanding of the genetic basis of ionome variation. Various biological mechanisms may drive the detection of these unique PC QTL. For example, the ionome has been shown to exhibit tissue-dependent, multi-element changes in response to nitrogen availability [24]. A unique PC QTL could be detected at a nitrogen metabolism gene if variation at that gene confers additive effects on multiple elements. Variation in genes involved in adaptive responses to drought stress, soil nutrient deficiencies, or toxic micronutrient levels, can result in covariation among several elements without particularly strong effects on a single element [1, 6, 25], making such genes only identifiable as QTL when working with multivariate traits.

The majority of molecularly identified ionomic mutants have multi-element effects. In particular, mutants in genes involved in Casparian strip function and associated root-based element flow, including *MYB36* [26], *ESB1* [27], and *LOTR1* [28], all display pleiotropic effects on multiple element accumulation in the leaves. In some cases, QTL affecting these traits might be detected using both single and multi-element approaches, as was the case with the chromosome 5 QTL we mapped for P, S, Fe, Mn, and Zn, as well as for PC1. However, if the changes to a suite of elements are small for individual elements or uncontrolled environmental conditions inflate the magnitude of error in measuring the genetic effects, a multi-ionomic trait may be a better fit for QTL detection. The fact that we detect both overlapping and unique sets of element and PC QTL suggests that single and multivariate approaches should be used in concert to avoid gaps in our understanding of element regulatory networks. The evidence suggests that some of the most interesting ionome homeostasis genes, including genes that are involved in environmental adaptation extending beyond the ionome, will be those best detected through multivariate methods.

In addition to being a tool for understanding the genetics of multi-element regulation, principal components also reflected environmental variation. An across-environment PCA of all lines was used to find variables that describe variation between lines among all 10 environments. The first two across-environment PCs capture most of the variation in the ionome across 10 different growouts, much of which is environmental. This can be seen in the ability of aPC1 and aPC2 to separate growouts by location and, in some cases, different years within a location. Thus, components from a PCA done across environments can capture the impact of environment on the ionome as a whole.

In our across-environment analysis, to account for different sets of IBM lines within environments, we tested an approach of projecting loadings from a PCA on a smaller set of lines onto the full data set. The similarity of the PJs and aPCs led us to conclude that the sampling effects of having different subsets of lines in each environment had little effect on the trait covariance estimation. This approach to validate aPCs may be useful in other studies that seek to connect data from disparate experiments and federate data collected by multiple laboratories. The method of deriving traits across environments using a small set of genotypic checks opens up the possibility of using multi-trait correlations across environments to permit very large scale GxE mapping experiments on data sets not initially intended for this purpose. Retrospective analysis of data, or further data generation from preexisting biological material present in both public and private spheres, is enabled by this approach. For example, multiple association panels have been constructed for trait mapping in maize. Typically, comparison of multi-trait correlations across different populations is inhibited by our inability to ensure the 1:1 correspondence of traits. By using the subset of lines common to all mapping populations to create a projection, comparable traits could be reflected onto to full datasets for comprehensive genetic evaluation and the loci detected in each panel could then be compared, as we have done here.

PCA on all environments is a way to find variation resulting from environmental factors that impact multiple elements, for example weather or soil variables. The weather data available to us for this study was limited to maximum and minimum temperature. We observed the strongest correlations for aPC1 and aPC2 during the third and fourth quarters of the growing season. Because seed filling occurs in the latter part of the season, temperature during this time could have a pronounced effect on seed elemental composition. However, the lack of striking correlations between environmental components and the projections and aPCs, environmental factors other than temperature must be the strongest factors. Information on soil properties provided insight into a potential driver of the environmental variability captured by aPC2, with a strong negative correlation between aPC2 and soil pH. Soil pH alters element availability in soil, and pH differences between locations should result in different kernel ionomes.

QTL were mapped to the aPCs that describe whole ionome variation across environments. These loci may encompass genes that pleiotropically affect the ionome in an environmentally-responsive manner. The correlation between aPC2 with pH as well as the finding of an aPC2 QTL for Mo exemplifies the possibility of using across-environment PCA to detect element homeostasis loci that respond to a particular environmental or soil variable and produce a multi-element phenotype. To the extent that these differences are adaptive, these alleles can contribute to local adaptation to soil environment and nutrient availability. The identification of aPC QTL indicates that the variation captured by aPCs has both environmental and genetic components. Our previous study using single element traits found extensive GxE in this dataset through formal tests, so it is not surprising that we see a large environmental component as well as genetic factors contributing to variation in the across-environment PCs. Experiments with more extensive weather and soil data, or carefully manipulated environmental contrasts, are needed to create models with additional covariates and precisely represent environmental impacts. This multivariate approach could be especially powerful in studies with extensive and consistent environmental variable recording, such as the “Genomes to Fields” Initiative, where specific environmental variables could be included in QTL models of multi-element GxE.

## Conclusions

Here we have shown that treating the ionome as an interrelated set of traits using PCA within environments can identify novel loci. PCA across environments allowed us to derive traits that described both environmental and genetic variation in the ionome.

## Methods

### Field Growth and Data Collection

#### Field growth and elemental profile analysis

Lines belonging to the Intermated B73 × Mo17 recombinant inbred (IBM) population [14] were grown in 10 different environments: Homestead, Florida in 2005 (220 lines) and 2006 (118 lines), West Lafayette, Indiana in 2009 (193 lines) and 2010 (168 lines), Clayton, North Carolina in 2006 (197 lines), Poplar Ridge, New York in 2005 (256 lines), 2006 (82 lines), and 2012 (168 lines), Columbia, Missouri in 2006 (97 lines), and Ukilima, South Africa in 2010 (87 lines). Elemental analysis was carried out in a standardized inductively coupled plasma mass spectrometry (ICP-MS) pipeline previously described in detail [15]. Analytical outlier removal and weight normalization was performed following data collection as described in our previous analysis of these data.

### Computational Analysis

#### Element correlation analysis

Within environments, 190 Pearson correlation coefficients were calculated, one for each pair of the 20 measured elements. To control for multiple tests, we applied a Bonferroni correction at an alpha level of 0.05. Given 190 possible combinations, correlations with a p-value below 0.05/190 = 0.00026 were regarded as significant.

#### Principal components analysis of ionome variation within environments

Elements prone to analytical error (B, Na, Al, As) were removed before to PC analysis, leaving 16 elements: Mg, P, S, K, Ca, Mn, Fe, Co, Ni, Cu, Zn, Se, Rb, Sr, Mo, and Cd. In an attempt to summarize the effects of genotype on covariance of ionomic components, a PCA was done using elemental data for each of the 10 environments separately. The *prcomp* function in R with scale = TRUE was used for PCA on elemental data to perform PCA on the line average element values in an environment. This function performs singular value decomposition on a scaled and centered version of the input data matrix, computing variances with the divisor *N*-1. 16 PCs were returned from each environment. After removal of PCs accounting for less than 2% of the variance, the 10 sets of PCs were used as traits in QTL analysis. Variance proportions and trait loadings for all PCs calculated across 10 environments are provided in S1 Table.

#### QTL Mapping: principal components

QTL mapping was done using stepwise forward-backward regression in R/qtl [29] as described previously for element phenotypes [15]. The mapping procedure was done for each environment separately, with PC line means for RILs in the given environment as phenotypes and RIL genotypes as input. The *stepwiseqtl* function was used to produce an additive QTL model for each PC, with the max number of QTL allowed for each trait set at 10. The 95^th^ percentile LOD score from 1000 *scanone* permutations was used as the penalty for addition of QTL. The QTL model was optimized using *refineqtl* for maximum likelihood estimation of QTL positions. The locations of the PC QTL detected in this study were compared to the single element QTL from our previous study. Loci were considered distinct if they were at least 25 cM away from any single element QTL detected in the environment in which the PC QTL was detected. This serves as a conservative control in order to minimize the mistaken assessment of novelty for QTL with small changes in peak position.

#### QTL by environment analysis: PCA across environments

The 16 most precisely measured elements were used for an additional principal components analysis. Again, the *prcomp* function in R with scale = TRUE was used for PCA on elemental data, however, all 16 element measurement values in all lines in all of the 10 environments were combined into one PCA. These PCs are referred to as across-environment PCs (aPCs). The first 7 aPCs explained 93% of the total covariation of these traits. A linear model was used to test the relationship of environmental parameters on these aPCs. All seven aPCs were also used for stepwise QTL mapping by the same method described above.

#### QTL by environment analysis: Projection-PCA across environments

The sets of lines grown in each our ten environments were drawn from the same population [14] but different subsets were grown and harvested in different environments. To achieve common multivariate summaries for all lines and growouts, we performed an alternative PCA using a smaller set of common lines. We then projected the loadings from this PCA onto the full dataset, as follows. First, a PCA was conducted on 16 lines common to six of the 10 environments (FL05, FL06, IN09, IN10, NY05, NY12). The loadings for each PC from this PCA were then used to calculate values from full set of lines across 10 environments to generate PCA projections (PJs). These derived values based on a common-line PCA were compared to previously described aPC values from the PCA done on all lines at once. Correlations between PJs and aPCs were computed to compare the outcomes of the two methods.

#### Weather and soil data collection and analysis

Weather data for FL05, FL06, IN09, IN10, NC06, NY05, NY06, and NY12 was downloaded from Climate Data Online (CDO), an archive provided by the National Climatic Data Center (NCDC) through the National Oceanic and Atmospheric Administration (http://www.ncdc.noaa.gov/cdo-web/). Data were not available for the South Africa growout. Daily summary data for each day of the growing season were tabulated from the weather station nearest to the field location. Weather stations used to obtain data for each location are indicated in S2 Table. Minimum temperature (in degrees Celsius) and maximum temperature (in degrees Celsius) were available in each location. With these variables, average minimum temperature, and maximum temperature were calculated across the 120-day growing season as well as for 30 day quarters. Growing degree days (GDD) were calculated for the entire season and quarterly using the formula GDD = ((Tmax + Tmin)/2) − 10.

Data describing soils from each location were obtained from the Web Soil Survey provided by the USDA Natural Resources Conservation Service (http://websoilsurvey.sc.egov.usda.gov/App/HomePage.htm). A representative area of interest was selected at the site of plant growth using longitude and latitude coordinates. When an area contained more than one soil type, a weighted average of measurements from all soil types was used. The data we downloaded from the Web Soil Survey were: pH, electrical conductivity (EC) (decisiemens per meter at 25 degrees C), available water capacity (AWC) (centimeters of water per centimeter of soil), available water supply (AWS) (centimeters), and calcium carbonate (CaCO3) content (percent of carbonates, by weight). Layer options were set to compute a weighted average of all soil layers.

The relationships between the seven experiment wide aPCs and the weather and soil variables were estimated by calculating Pearson correlation coefficients for the pairwise relationships. Correlations were also calculated between average element values and soil and weather variables in each environment.

## Acknowledgements

The authors would especially like to thank our field collaborators Sherry Flint-Garcia, Peter Balint-Kurti, Torbert Rocheford, Jonathan Lynch, and Robert Snyder for their dedicated efforts to provide the seeds analyzed for this project. This work was supported by funding from the National Science Foundation (IOS-1126950, IOS-1450341), the USDA Agricultural Research Service (5070-21000-039-00D). AA was a recipient of a Danforth Plant Science Fellowship from the Donald Danforth Plant Science Center.

## Supporting Information

**S1 Fig. Variances of Principal Components from PCA within 10 Environments.** Eigenvalues (amount of variation explained) for each PC are shown on the y-axis. Lines are colored by environment.

**S2 Fig. Loadings of Principal Components from Different Environments.** Loadings for each element are plotted for PCs from different environments. Loadings of PCs plotted on the same graph are correlated as indicated. PCs shown in (A), (B), and (C) all have a QTL coinciding with Mo QTL on chromosome 1. PCs shown in (D) have a QTL coinciding with Cd QTL on chromosome 2. PCs shown in (E) have a QTL coinciding with Ni QTL on chromosome 9.

**S3 Fig. Variances of Principal Components from PCA on Lines from all Environments.** Eigenvalues (amount of variation explained) for each aPC are shown on the y-axis.

**S4 Fig. aPC1 and aPC2 Loadings Biplot.** PCA plots showing aPC1 and aPC2 loadings. Variance explained for each PC is indicated along axes.

**S1 Table. PC Variance Proportions and Loadings Across 10 Environments.**

**S2 Table. Weather Station Locations.**

## References

1. Baxter IR, Vitek O, Lahner B, Muthukumar B, Borghi M, Morrissey J, et al. The leaf ionome as a multivariable system to detect a plant’s physiological status. Proceedings of the National Academy of Sciences. 2008;105: 12081–12086.

2. Korshunova YO, Eide D, Clark WG, Guerinot ML, Pakrasi HB. The IRT1 protein from Arabidopsis thaliana is a metal transporter with a broad substrate range. Plant molecular biology. 1999;40: 37–44.

3. Broadley MR, Hammond JP, King GJ, Astley D, Bowen HC, Meacham MC, et al. Shoot calcium and magnesium concentrations differ between subtaxa, are highly heritable, and associate with potentially pleiotropic loci in Brassica oleracea. Plant Physiol. 2008;146: 1707–1720.

4. Buescher E, Achberger T, Amusan I, Giannini A, Ochsenfeld C, Rus A, et al. Natural genetic variation in selected populations of Arabidopsis thaliana is associated with ionomic differences. PLoS One. 2010;5: e11081.

5. Baxter IR, Gustin JL, Settles AM, Hoekenga OA. Ionomic characterization of maize kernels in the intermated B73 × Mo17 population. Crop Science. 2013;53: 208–220.

6. Baxter I. Ionomics: studying the social network of mineral nutrients. Current Opinion in Plant Biology. 2009;12: 381–386.

7. Burton AL, Johnson J, Foerster J, Hanlon MT, Kaeppler SM, Lynch JP, et al. QTL mapping and phenotypic variation of root anatomical traits in maize (Zea mays L.). Theoretical and Applied Genetics. 2015;128: 93–106.

8. Bouchet S, Bertin P, Presterl T, Jamin P, Coubriche D, Gouesnard B, et al. Association mapping for phenology and plant architecture in maize shows higher power for developmental traits compared with growth influenced traits. Heredity. 2017; 118: 249–259.

9. Frey FP, Presterl T, Lecoq P, Orlik A, Stich B. First steps to understand heat tolerance of temperate maize at adult stage: identification of QTL across multiple environments with connected segregating populations. Theoretical and Applied Genetics. 2016;129: 945–961.

10. Topp CN, Iyer-Pascuzzi AS, Anderson JT, Lee C-R, Zurek PR, Symonova O, et al. 3D phenotyping and quantitative trait locus mapping identify core regions of the rice genome controlling root architecture. Proceedings of the National Academy of Sciences. 2013; 110: E1695–E1704.

11. Liu Z, Garcia A, McMullen MD, Flint-Garcia SA. Genetic Analysis of Kernel Traits in Maize-Teosinte Introgression Populations. G3: Genes Genomes Genetics. 2016;6: 2523.

12. Choe E, Rocheford TR. Genetic and QTL analysis of pericarp thickness and ear architecture traits of Korean waxy corn germplasm. Euphytica. 2012;183: 243–260.

13. Zhang N, Gibon Y, Gur A, Chen C, Lepak N, Hohne M, et al. Fine Quantitative Trait Loci Mapping of Carbon and Nitrogen Metabolism Enzyme Activities and Seedling Biomass in the Intermated Maize IBM Mapping Population. Plant Physiology. 2010; 154: 1753–1765.

14. Lee M, Sharopova N, Beavis WD, Grant D, Katt M, Blair D, et al. Expanding the genetic map of maize with the intermated B73 × Mo17 (IBM) population. Plant molecular biology. 2002;48: 453–461.

15. Asaro A, Ziegler G, Ziyomo C, Hoekenga OA, Dilkes BP, Baxter I. The Interaction of Genotype and Environment Determines Variation in the Maize Kernel Ionome. G3: Genes Genomes Genetics. 2016;6: 4175–4183.

16. Poormohammad Kiani S, Trontin C, Andreatta M, Simon M, Robert T, Salt DE, et al. Allelic Heterogeneity and Trade-Off Shape Natural Variation for Response to Soil Micronutrient. PLOS Genetics. 2012;8: e1002814.

17. Barberon M, Vermeer J, De Bellis D, Wang P, Naseer S, Andersen T, et al. Adaptation of Root Function by Nutrient-Induced Plasticity of Endodermal Differentiation. Cell. 2016;164: 447–459.

18. Chao DY, Gable K, Chen M, Baxter I, Dietrich CR, Cahoon EB, et al. Sphingolipids in the Root Play an Important Role in Regulating the Leaf Ionome in Arabidopsis thaliana. Plant Cell. 2011;23: 1061–1081.

19. Lopez-Arredondo DL, Leyva-Gonzolez MA, Gonzolez-Morales SI, Lopez-Bucio J, Herrera-Estrella L. Phosphate nutrition: improving low-phosphate tolerance in crops. Annual Review of Plant Biology. 2014;65: 95–123.

20. Lin Y-F, Liang H-M, Yang S-Y, Boch A, Clemens S, Chen C-C, et al. Arabidopsis IRT3 is a zinc-regulated and plasma membrane localized zinc/iron transporter. New Phytologist. 2009;182: 392–404.

21. Mulder EG. Importance of molybdenum in the nitrogen metabolism of microorganisms and higher plants. Plant and Soil. 1948; 1: 94–119.

22. Mulder EG, Gerretsen FC. Soil manganese in relation to plant growth. Adv Agron. 1952;4: 221–277.

23. Millikan CR. Antagonism between molybdenum and certain heavy metals in plant nutrition. Nature. 1948; 161: 528.

24. Chu Q, Watanabe T, Shinano T, Nakamura T, Oka N, Osaki M, et al. The dynamic state of the ionome in roots, nodules, and shoots of soybean under different nitrogen status and at different growth stages. 2016;17: 488–498.

25. Baxter I, Dilkes BP. Elemental Profiles Reflect Plant Adaptations to the Environment. Science. 2012;336: 1661–1663.

26. Kamiya T, Borghi M, Wang P, Danku JMC, Kalmbach L, Hosmani PS, et al. The MYB36 transcription factor orchestrates Casparian strip formation. Proceedings of the National Academy of Sciences. 2015;112: 10533–10538.

27. Baxter I, Hosmani PS, Rus A, Lahner B, Borevitz JO, Muthukumar B, et al. Root suberin forms an extracellular barrier that affects water relations and mineral nutrition in Arabidopsis. PLoS Genet. 2009;5: e1000492.

28. Li B, Kamiya T, Kalmbach L, Yamagami M, Yamaguchi K, Shigenobu S, et al. Role of LOTR1 in Nutrient Transport through Organization of Spatial Distribution of Root Endodermal Barriers. Current Biology. 2017;27: 758–765.

29. Broman KW, Speed TP. A model selection approach for the identification of quantitative trait loci in experimental crosses. Journal of the Royal Statistical Society: Series B (Statistical Methodology). 2002;64: 641–656.

